# A comparative meta and *in silico* analysis of differentially expressed genes and proteins in canine and human bladder cancer

**DOI:** 10.1101/2020.05.04.039719

**Authors:** Victoria Vitti Gambim, Renee Laufer-Amorim, Ricardo Fonseca Alves, Valeria Grieco, Carlos Eduardo Fonseca-Alves

## Abstract

Canine and human bladder cancer present several similar anatomical, morphological and molecular characteristics and dogs can be considered a model for human bladder cancer. However, the veterinary literature lacks information regarding cross validation analysis between human and canine large-scale data. Therefore, this research aimed to perform a meta-analysis of the previous canine literature on bladder cancer, identifying genes and protein previously evaluated in these studies. Besides that, we also performed a cross validation of the canine transcriptome data and the human data from The Cancer Genome Atlas (TCGA) to identify potential markers for both species. It was performed a meta-analysis using the following indexing terms “bladder” AND “carcinoma” AND “dog” in different international databases and 385 manuscripts were identified in our initial search. Then, several inclusion criteria were applied and only 25 studies met these criteria. Among these studies, five presented transcriptome data and 20 evaluated only isolated genes or proteins.

Regarding the studies involving isolated protein analysis, HER-2 protein was the most studied (3/20), followed by TAG-72 (2/20), COX-2 (2/2), Survivin (2/2) and CK7 (2/2). Regarding the cross-validation analysis of human and canine transcriptome data, we identified 35 deregulated genes, including *ERBB2, TP53, EGFR* and *E2F2*. Our results demonstrated that the previous canine literature on bladder cancer was focused on the evaluation of isolated markers with no association with patient’s survival. Besides that, the lack of information regarding tumor muscle-invasion can be considered an important limitation when comparing human and canine bladder tumors. Our in-silico analysis involving canine and human transcriptome data provided several genes with potential to be markers for both human and canine bladder tumors and these genes should be considered for future studies on canine bladder cancer.

## 1 Introduction

Transitional cell carcinoma (TCC), also called urothelial carcinoma, is the most common bladder cancer in both humans and dogs, sharing clinical, pathological and molecular alterations (Dhawan et al., 2018, Maeda et al., 2018, Ramsey et al., 2017). In the United States, it is expected 81.400 new cases and 17.980 bladder cancer-related deaths (Siegel et al., 2020). The last global cancer statistics (GLOBOCAN) revealed 549.393 new cases and 199.922 bladder cancer-related deaths (Bray et al. 2018). In dogs, urothelial carcinoma is the most common malignant tumor in canine bladder, representing 1% of all neoplasms that affect dogs (Patrick et al., 2006). In humans, TCC is a tumor associated with several factors such as cigarette smoking, occupational exposure (Myasaki and Nishiyama, 2017), arsenic and aromatic compounds (Cha et al., 2018). In household dogs, a case-control study was previously performed to correlate cigarette smoke, obesity, use of topical insecticides and chemicals used at home with canine bladder cancer development (Glickman et al. 1989). These authors founded high risk of bladder cancer development in obese dogs and dogs that used topical insecticides (Glickman et al. 1989). Since dogs and humans shares the same environment, dogs could be considered a sentinel (Knapp et al., 2014).

Canine and human TCC are usually a local infiltrative cancer, that can extend to the entire bladder, including submucosa and muscular layers (Grzegółkowski et al., 2016, Andrade et al., 2004). Besides that, human bladder carcinoma can invade adjacent tissues and organs such as ureter, prostatic urethra and prostate gland (Hernández-Fernández et al., 2016). Usually, human bladder carcinoma are superficial tumors (70% of the cases) and are classified as a non-muscle invasive bladder carcinoma (NMIBC) (Grzegółkowski et al., 2016). Since NMIBC presents better prognosis, the muscle invasive bladder carcinoma is considered a challenge (Grzegółkowski et al., 2016, Hernández-Fernández et al., 2016). Thus, in the human literature, the muscle invasive bladder carcinoma is focus of recent studies. In dogs, the infiltration is not standardized, since in several cases tissue samples comes from cystoscopy (Dhawan et al. 2018, Dhawan et al. 2015).

The molecular phenotype of human bladder cancer is widely studied, and some genomic subtyping was previously proposed (Jalanko et al. 2020, Inamura et al. 2018). The Cancer Genome Atlas (TCGA) database revealed 64 significantly muted genes in human TCC, including p53/cell cycle, DNA repair, PI3K/AKT, and chromatin modifications and regulation. Besides that, there is a subclassification for human muscle invasive bladder cancer, luminal-papillary, luminal-infiltrated, luminal, basal-squamous and neuronal subtypes (Cancer Genome Atlas Research Network, 2014). On the other hand, there are few papers performing molecular characterization of canine bladder cancer and it is considered a promising area and the molecular characterization of canine bladder carcinoma can provide a valuable information regarding this tumor biological behavior (Knapp et al., 2020).

In dogs, some recent studies performing transcriptome analysis revealed several important molecular findings, such as differentiation of canine bladder cancer in molecular subtypes according to *BRAFV595E* somatic mutation (*BRAFV600E* in humans) (Parker et al. 2020), identification therapeutic targets (*PTGER2, ERBB2, CCND1, VEGF*, and *EGFR*) and separation in basal and luminal subtypes, as for human bladder cancers, in order to compare their different muscular invasive potential (Dhawan et al., 2018). Therefore, studies performing comparative molecular comparisons between human and canine bladder carcinomas can provide a unique opportunity to study this cancer subtype in both species. In this scenario, this manuscript aimed to perform a literature meta-analysis and extract all information regarding gene and protein expression in canine bladder cancer and perform an *in silico* analysis to identify common gene alterations among dogs and human to select candidates for future studies regarding prognosis or treatment.

## 2 Material and Methods

### 2.1 Study design

The study design is summarized in figure 1. We divided the study methods in three steps: 1) metanalysis of the previous literature aiming to identify deregulated genes and proteins in canine bladder cancer, 2) *In silico* analysis of deregulated gene and proteins to identify prognostic and predictive marker in canine bladder cancer and 3) Were selected five previous studies with transcriptome data and we extracted common gene information from these studies and validated with The Cancer Genome Atlas (TCGA) data.

**Figure 1.**
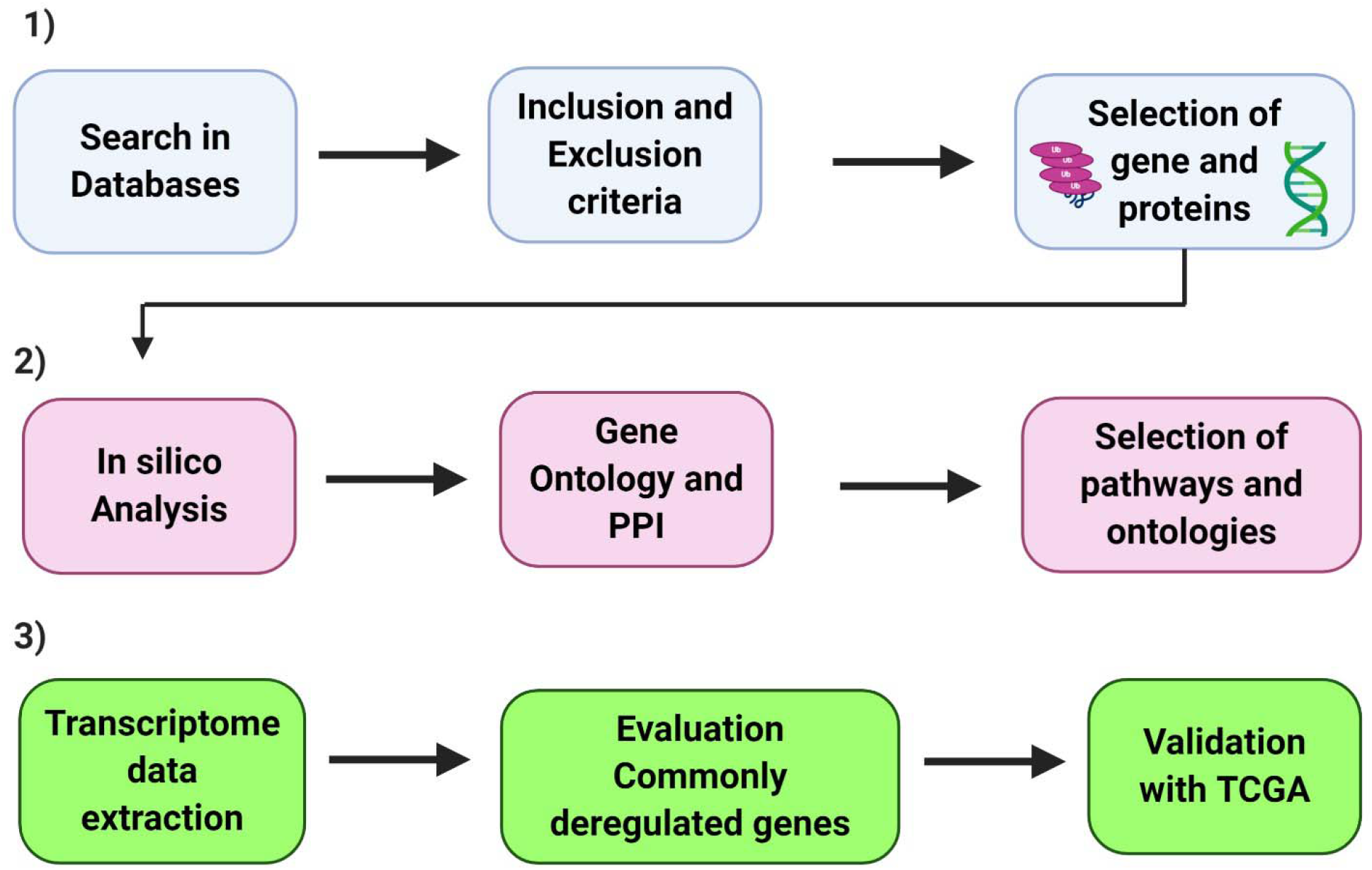
Schematic representation of the manuscript design. 1) we searched in different databases manuscripts evaluating proteins or genes in canine bladder carcinomas. Between the identified manuscripts, we applied different inclusion and exclusion criteria to extract the gene and protein information from these manuscripts. Therefore, we performed two different analysis based on the manuscript data. 2) manuscripts with information regarding isolated genes or proteins were evaluated together. We extracted the name of the gene or protein and the p value to perform ontology and protein-protein interaction analysis. In the end of this analysis, we identified different pathways and ontology process related to the previous published data. 3) the manuscripts with large-scale transcriptome data were evaluated together in a separated analysis. We identified genes commonly deregulated among studies and cross-validated this data with The Cancer Genome Atlas (TCGA). The diagram was generated using BioRender (https://app.biorender.com/).

### 2.2 Meta-analysis

In order to identify previous published papers to include in our meta-analysis, we performed a literature search in the following databases: *PubMed, MEDLINE* and *Scielo* using the following indexing terms “bladder” AND “carcinoma” AND “dog” with no restriction regarding the year of publication. Then, we reviewed the reference section of the selected manuscripts and performed a manual search in the most relevant journals with veterinary oncology background to ensure we included the highest number of available manuscripts.

Afterwards, we selected manuscripts by title and abstract, including scientific articles that evaluated gene or proteins in canine bladder carcinomas. In this step, we excluded review manuscripts, case reports and retrospective studies including only survival analysis. Then, we analyzed each included manuscript and selected scientific papers that evaluate gene or proteins in canine bladder samples, comparing with normal bladder tissues. In this step, we excluded manuscript using only cell lines, manuscripts that compared bladder carcinomas to other conditions such as cystitis and manuscripts evaluating only bladder carcinomas, with no comparison to normal bladder tissue. Our first evaluation was performed on December 12^th^, 2019 and it was last updated in April 2^nd^, 2020.

From the selected manuscripts, we retrieved information’s regarding each deregulated gene or protein, the “p-value” for each gene or protein (comparison between bladder cancer and normal bladder) and survival.

### 2.3 *In silico* analysis

The *in-silico* analysis of each evaluated gene and protein was performed using online free-evaluable tools. In the first step, we selected only the deregulated genes and evaluated by *STRING* (https://string-db.org/) to translate into each respective protein. Then, we used only proteins for the subsequent analysis. We opted to evaluate only proteins in our *in-silico* study due the facility to use proteins as prognostic and predictive markers.

The deregulated proteins were evaluated grouped (upregulated and downregulated) or independently (upregulated or downregulated) using the online Search Tool for the Retrieval of Interacting Genes - STRING (https://string-db.org/) to generate protein-protein interaction (PPI) networks. We considered only STRING interactions of high confidence (0.700) and we hid the disconnected nodes for a better visualization. The considered interaction to generate PPI networks were co-expression, cooccurrence, databases and neighborhood interactions.

### 2.4 Gene ontology

The gene ontology (GO) analysis was performed to facilitate the comprehension of results among different species. The selected proteins were analyzed using Enrichr (https://amp.pharm.mssm.edu/Enrichr/) aiming to sample enrichment and GO evaluation. The analyzed information were retrieved from Enrichr and submitted to Revigo (http://revigo.irb.hr/) to organize and visualize the enriched GO terms. We opted to use the three GO categories: biological process, cellular component and molecular function. However, since biological process and molecular function terms have a higher chance to provided prognostic and predictive information, we focused on these two categories.

### 2.5 Transcriptome data retrieved from the previous literature

We selected three previous studies that evaluated the transcriptome (RNA-seq) of canine bladder carcinoma, with the data sets available online at NCBI short-read archive (SRA) under BioProject ID PRJNA559406 (Parker et al., 2020), GEO database (ref: GSE24152) (Dhawan et al., 2018) or available at the DDBJ Sequenced Read Archive repository (http://trace.ddbj.nig.ac.jp/dra/index_e.html) with accession number DRA005844). Besides that, we also included one manuscript evaluating the transcriptome of canine bladder cancer using microarray data (Dhawan et al., 2015) and one manuscript that performed both mRNA-seq and exome-seq (Ramsey et al. 2017). For both studies (Dhawan et al 2015 and Ramsey et al. 2017) mRNA data were obtained in supplementary information.

The genes differentially expressed between TCC and normal bladder were selected following the criteria p< 0.05, 2.5-fold change or higher and −2.5-fold change or lower. Venn diagram was performed using an online tool (http://www.interactivenn.net/) (Herbele et al., 2015). Furthermore, the commonly differentially expressed genes among the five studies were validated using the 344 bladder carcinomas from TCGA database.

### 2.6 The Cancer Genome Atlas (TCGA) and The Cancer Proteome Atlas (TCPA) crossvalidation

Due the lack of a veterinary database with deposited information regarding survival of canine patient with bladder carcinoma, we selected the most relevant proteins and validated using 344 human samples from patients with bladder carcinoma from TCGA (https://www.cancer.gov/tcga). Then, the cross-validated proteins were evaluated by TCPA (https://tcpaportal.org/tcpa/index.html) (Juan et al., 2017; Li et al., 2013). We considered 5% interval of confidence or protein integrations with p-value lower than 0.05.

Besides that, we performed two different analysis using the TCPA. First, we used “visualization” tool to perform a global analysis to evaluate interaction among genes, including negative and positive correlations. Thus, we selected genes and pathways of human bladder cancer differentially expressed as possible markers to be used in canine bladder cancer. In the second analysis, we evaluated the overall survival of the 344 human bladder cancer patients, according to protein expression level (high versus low). In this analysis, we selected all genes in human bladder cancer with prognostic value. The Kaplan Meier curves were generated using the TCPA online tool “individual cancer analysis (https://tcpaportal.org/tcpa/analysis.html) (Jun et al., 2013, Jun et al., 2017).

## 3 Results

### 3.1 Meta-analysis

385 manuscripts were identified in the initial search. Then, according to inclusion criteria, after reading the title and abstract we excluded 329 manuscripts and after reading the full manuscript, 25 of them met our inclusion criteria (Figure 2). Afterwards, we divided the selected manuscript in two categories, being manuscript with global transcriptome analysis (N=5) and manuscript with reported isolated genes or proteins (N=20). A complete list with the selected manuscripts can be found in Table 1.

**Figure 2.**
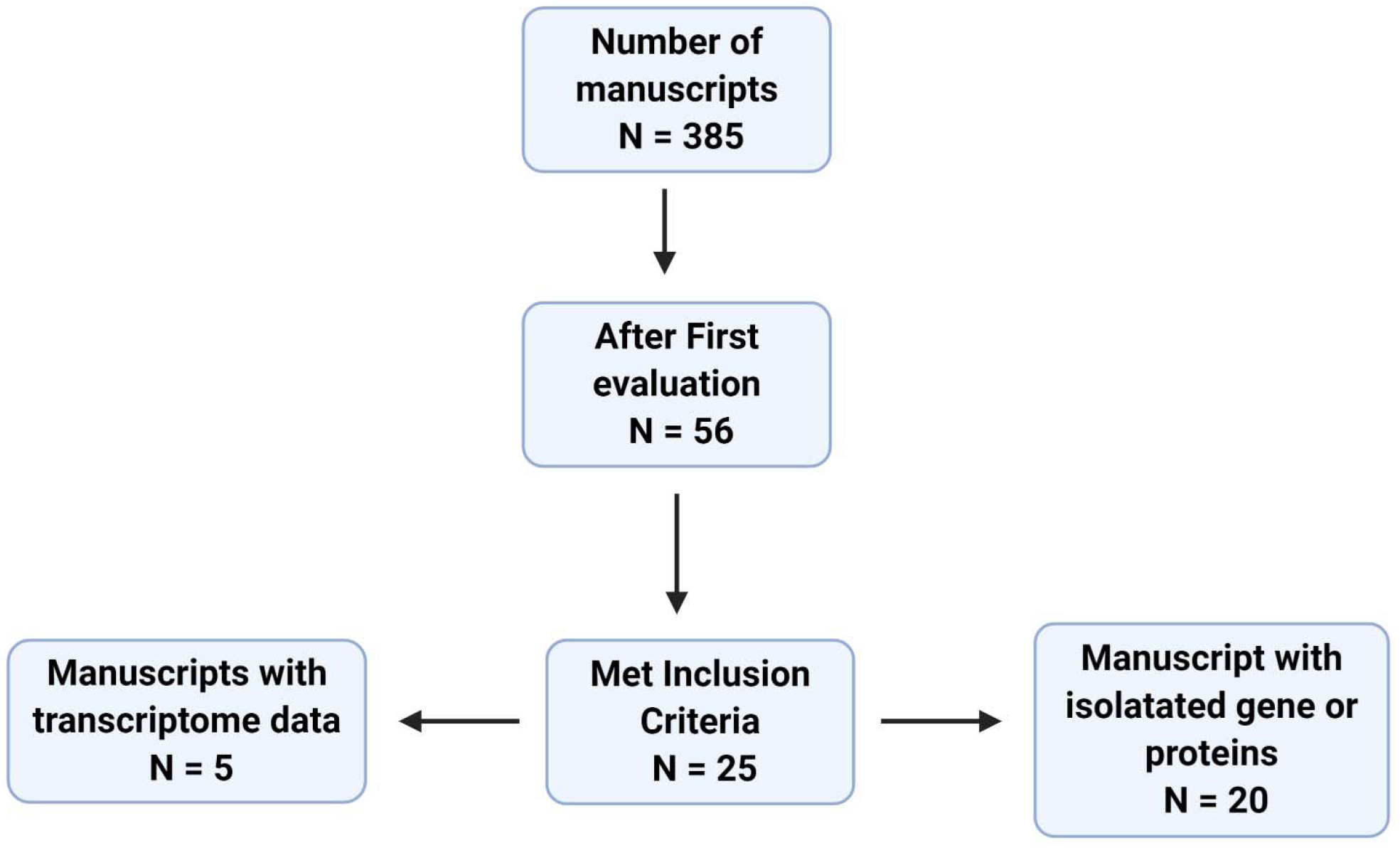
Flowchart of the evaluated manuscripts. In the first search, it was identified 385 manuscript and after applying all inclusion and exclusion criteria we identified five manuscripts with large-scale transcriptome data and 20 manuscript with isolated genes or proteins. The diagram was generated using BioRender (https://app.biorender.com/).

**Table 1.**
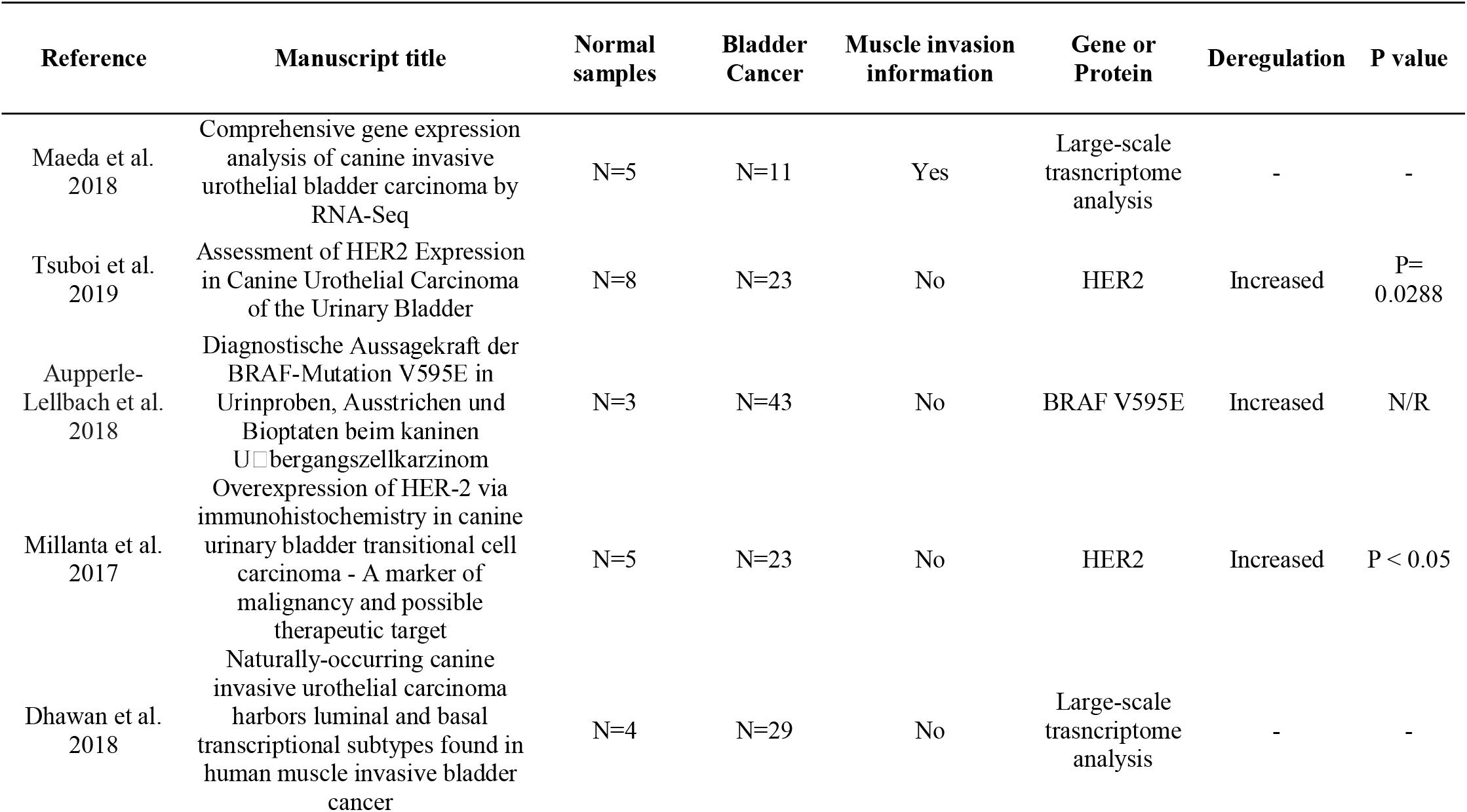
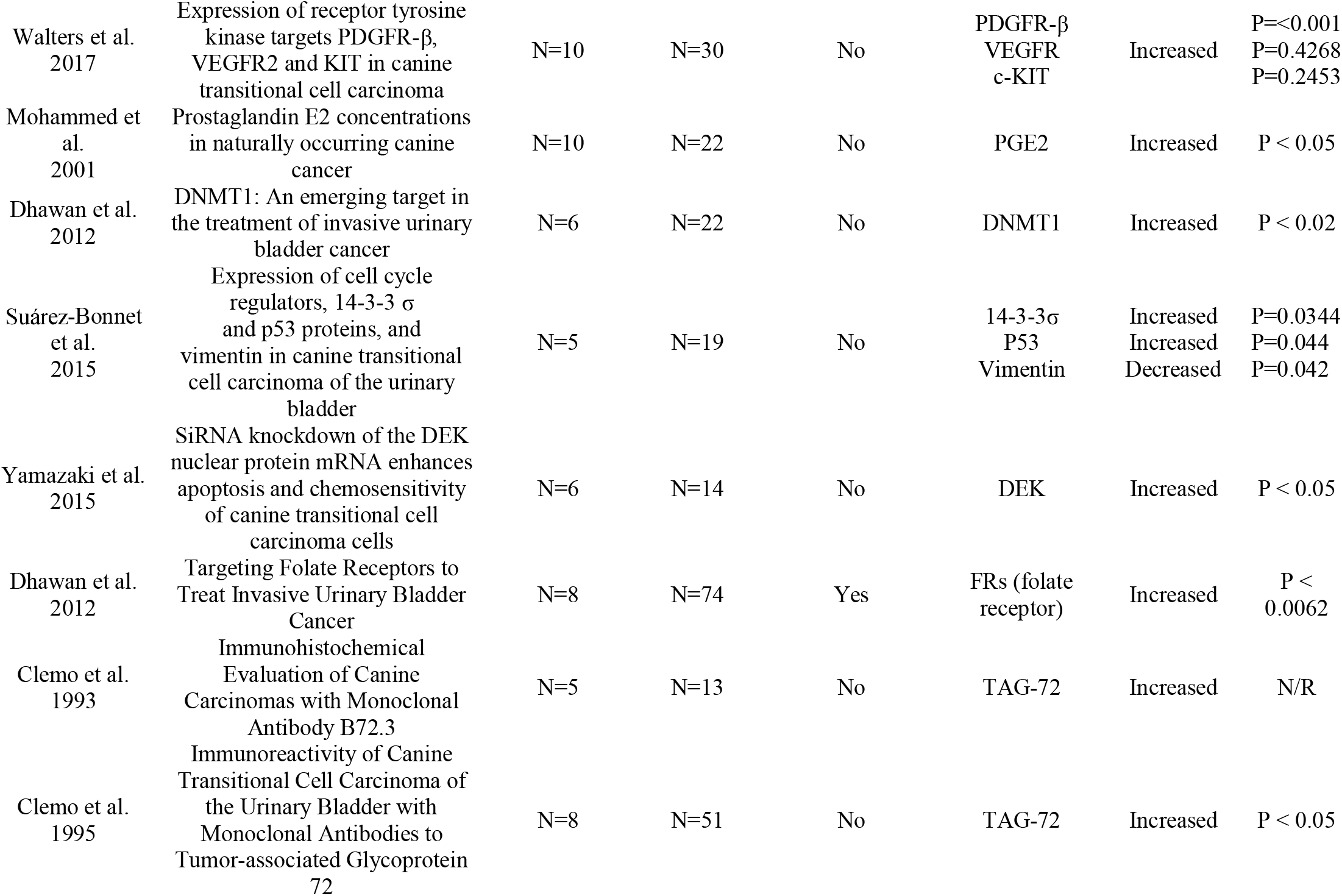
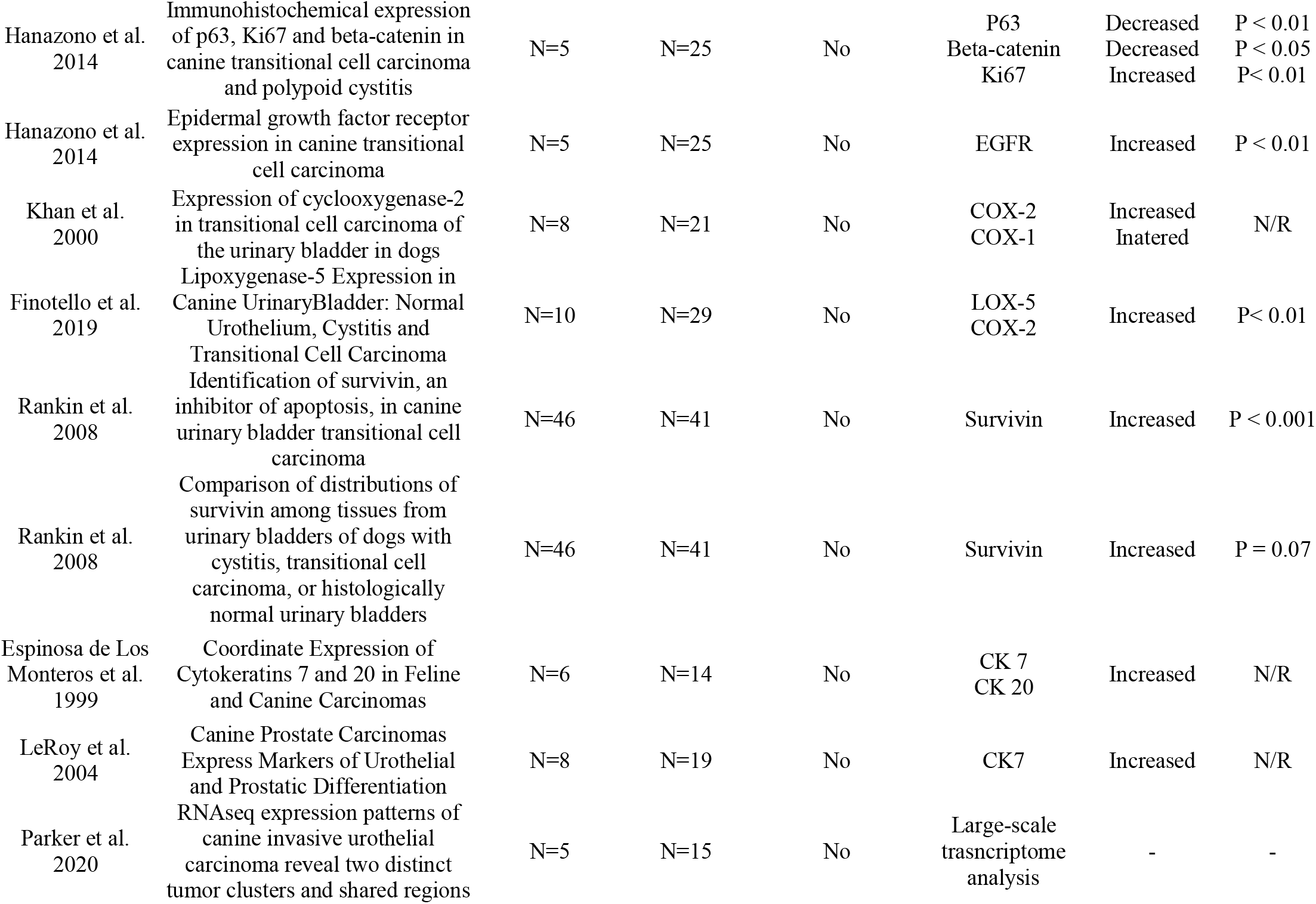
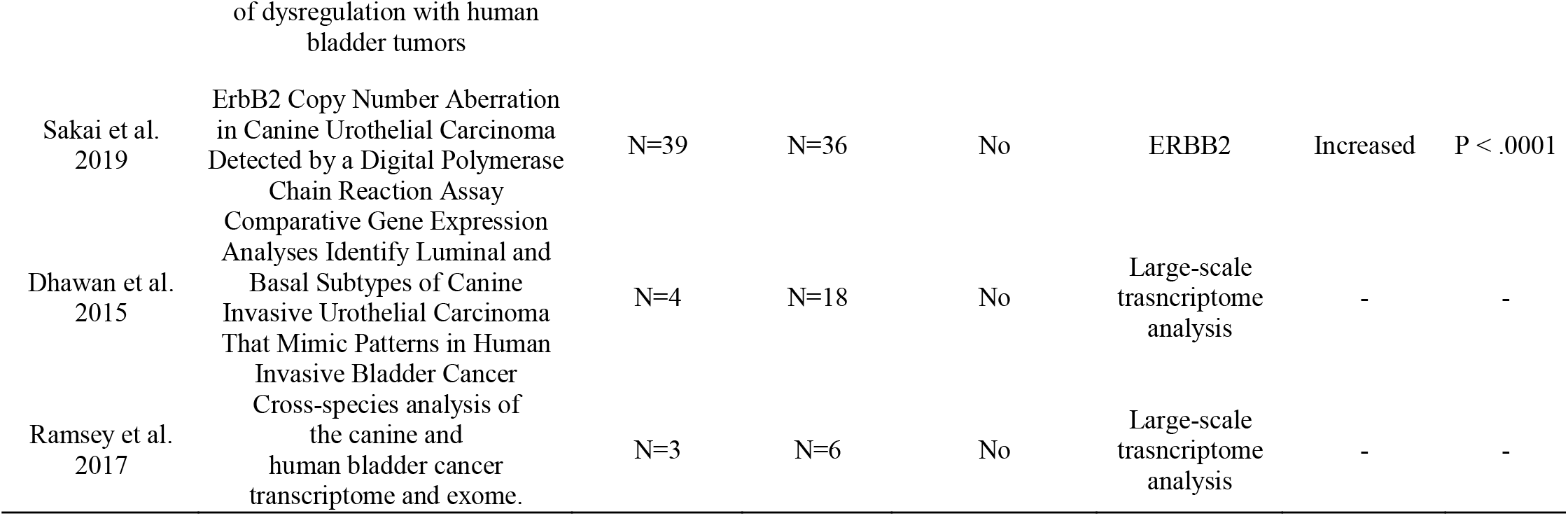
Manuscripts (N=25) that met our inclusion criteria and were used in the research of canine bladder carcinoma.

Regarding the studies involving isolated protein analysis, HER-2 was the most studied protein (3/20), followed by TAG-72 (2/20), COX-2 (2/2), Survivin (2/2) and CK7 (2/2). The remaining proteins were evaluated in only one previous study each (Table 1). In the protein-protein interaction analysis, we identified one interaction network among proteins, being P53 the protein with the highest number of interactions. After enrichment analysis using Enrichr, we evaluated the most common ontology process associated with the previous published protein and we identified several processes related to tyrosine kinase regulation, cell communication and signaling and MAPK pathway (Supplementary Figure 1 and Supplementary Table 1).

### 3.2 In silico analysis of canine transcriptome data

In our meta-analysis, we identified five previous studies containing transcriptome data and in the most recent study (Parker et al., 2020), they cross validated their finding with other three published manuscripts (Dhawan et al., 2018, Maeda et al. 2018, Dhawan et al., 2015). Thus, we opted to analyze the transcriptome data from these five previous studies and cross validate with TCGA data. In our cross-validation analysis, we identified 61 deregulated genes (Figure 3), including CD55, IL17B, EGFR, CDH17 and CDH26. Moreover, we performed a PPI analysis among these genes and demonstrated a high interaction among them, including VEGFA, EGFR, TNF and CCND1 as central genes in the interaction network (Figure 3).

We identified 35 deregulated genes in cross-validation among the five veterinary studies and the TCGA data (Table 2).

**Figure 3.**
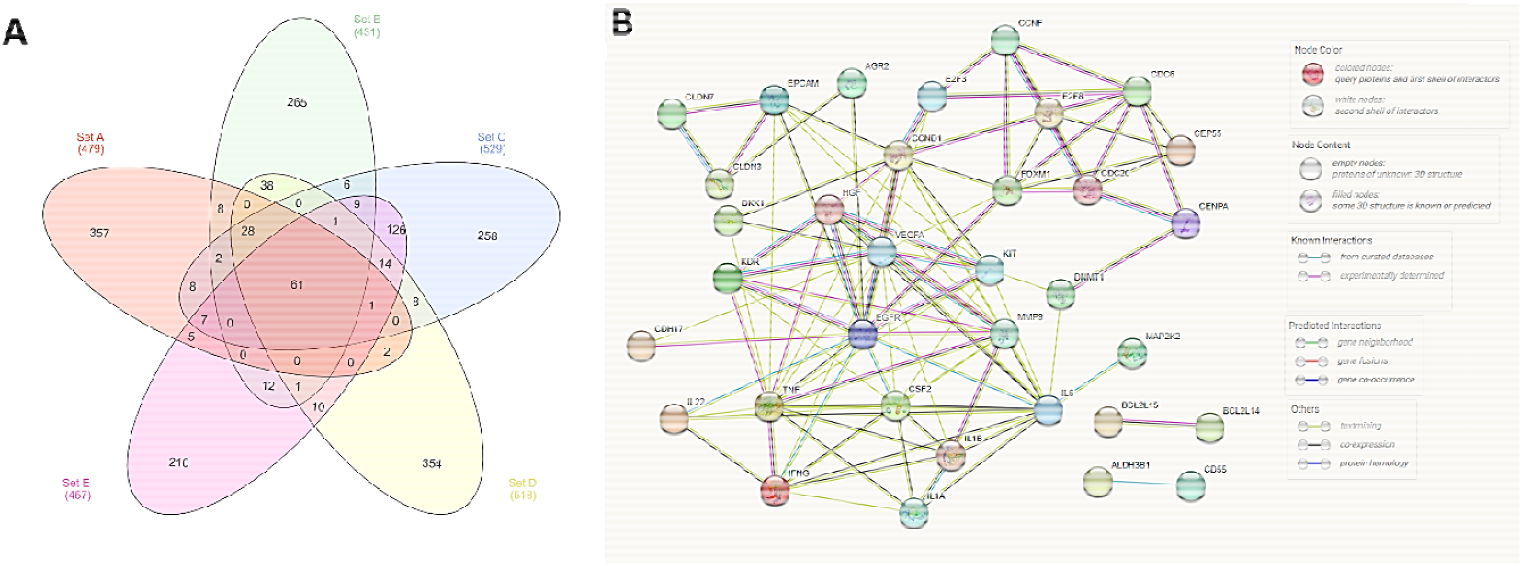
Transcriptome analysis of the five previous published studies in canine bladder transitional cell carcinoma. A: Venn diagram demonstrating the number of commonly deregulated genes among the five studies, including 61 genes commonly deregulated. B: Protein-protein interaction (PPI) network of the 61 deregulated genes. It is possible to observe several interactions among genes, including *EGFR* and *VEGFA* with high interaction networks. The Veen diagram was generated online (http://www.interactivenn.net/) and PPI was generated using STRING (https://string-db.org/).

**Table 2.** Deregulated genes after the cross-validation of the five veterinary studies with transcriptome data and the 344 human samples from The Cancer Genoma Atlas (TCGA).

| Gene/Protein | Chromosome | Gene ID | Regulation | Function |
| --- | --- | --- | --- | --- |
| BRAF V595E | 16 | 475526 | Upregulated | proto-oncogene, serine/threonine kinase |
| PTGER2 | 8 | 403797 | Upregulated | prostaglandin E receptor 2 |
| ERBB2 / HER-2 | 9 | 403883 | Upregulated | receptor tyrosine kinase 2 |
| CHST4 | 5 | 489722 | Upregulated | carbohydrate sulfotransferase 4 |
| PIGR | 7 | 474357 | Upregulated | polymeric immunoglobulin receptor |
| S100A14 | 7 | 612322 | Upregulated | calcium binding protein A14 |
| ADGRF1 | 12 | 474927 | Upregulated | adhesion G protein-coupled receptor F1 |
| AGR2 | 14 | 482333 | Upregulated | anterior gradient 2, protein disulphide isomerase |
| Irgm 1 | 11 | 606863 | Downregulated | immunity-related GTPase family M protein-like |
| TP53 | 5 | 403869 | Downregulated | tumor protein |
| ZFP36 | 1 | 484510 | Downregulated | ring finger protein |
| E2F2 | 2 | 100855664 | Upregulated | transcription factor 2 |
| IFI16 | 38 | 488622 | Upregulated | interferon-activable protein 203 |
| EGFR | 18 | 404306 | Upregulated | epidermal growth factor receptor |
| ERK1/2 | 26 | 477575 | Upregulated | mitogen-activated protein kinase 1 |
| NfκB-RelA | 12 | 481711 | Upregulated | NFKB inhibitor like 1 |
| MAP2K3 | 5 | 489547 | Upregulated | mitogen-activated protein kinase kinase 3 |
| IL1A | 17 | 403782 | Upregulated | interleukin 1 alpha |
| RABL6 | 9 | 480675 | Upregulated | RAS oncogene family like 6 |
| CCND1 | 18 | 449028 | Upregulated | cyclin D1 |
| FOXM1 | 27 | 486743 | Upregulated | forkhead box M1 |
| E2F3 | 35 | 488239 | Upregulated | transcription factor 3 |
| FOXO1 | 6 | 609116 | Upregulated | RNA binding fox-1 homolog 1 |
| IL6 | 14 | 403985 | Upregulated | interleukin 6 |
| IL1B | 17 | 403974 | Upregulated | interleukin 1 beta |
| CSF2 | 11 | 403923 | Upregulated | colony stimulating factor 2 |
| TNF | 12 | 403922 | Upregulated | tumor necrosis factor |
| IFNG | 10 | 403801 | Upregulated | interferon gamma |
| IL22 | 10 | 481153 | Upregulated | interleukin 22 |
| HGF | 18 | 403441 | Upregulated | hepatocyte growth factor |
| VEGF | 12 | 403802 | Upregulated | vascular endothelial growth factor A |
| PDGFR-B | 4 | 442985 | Upregulated | platelet derived growth factor receptor beta |
| VEGFR2 | 13 | 482154 | Upregulated | kinase insert domain receptor |
| DNMT1 | 20 | 476715 | Upregulated | DNA methyltransferase 1 |
| SFN | 2 | 487351 | Upregulated | Stratifin |

### 3.3 Analysis of the 344 human bladder cancer

In the analysis of the human bladder cancer samples, we identified several protein interactions, being positive or negative correlation (Figure 4). Regarding the PPI, it was possible to observe a high number of proteins from tyrosine kinase family, such as EGFR, ERK2, ERBB2 and BRAF. Besides that, we identified 28 proteins with prognostic value in human bladder cancer (Table 3). Among these proteins, only two were previously studied in canine bladder cancer (2/26). The top six proteins with prognostic value were Annexin1, TAZ, SF2, SRC, ARID1A and GATA3 (Figure 5).

**Figure 4.**
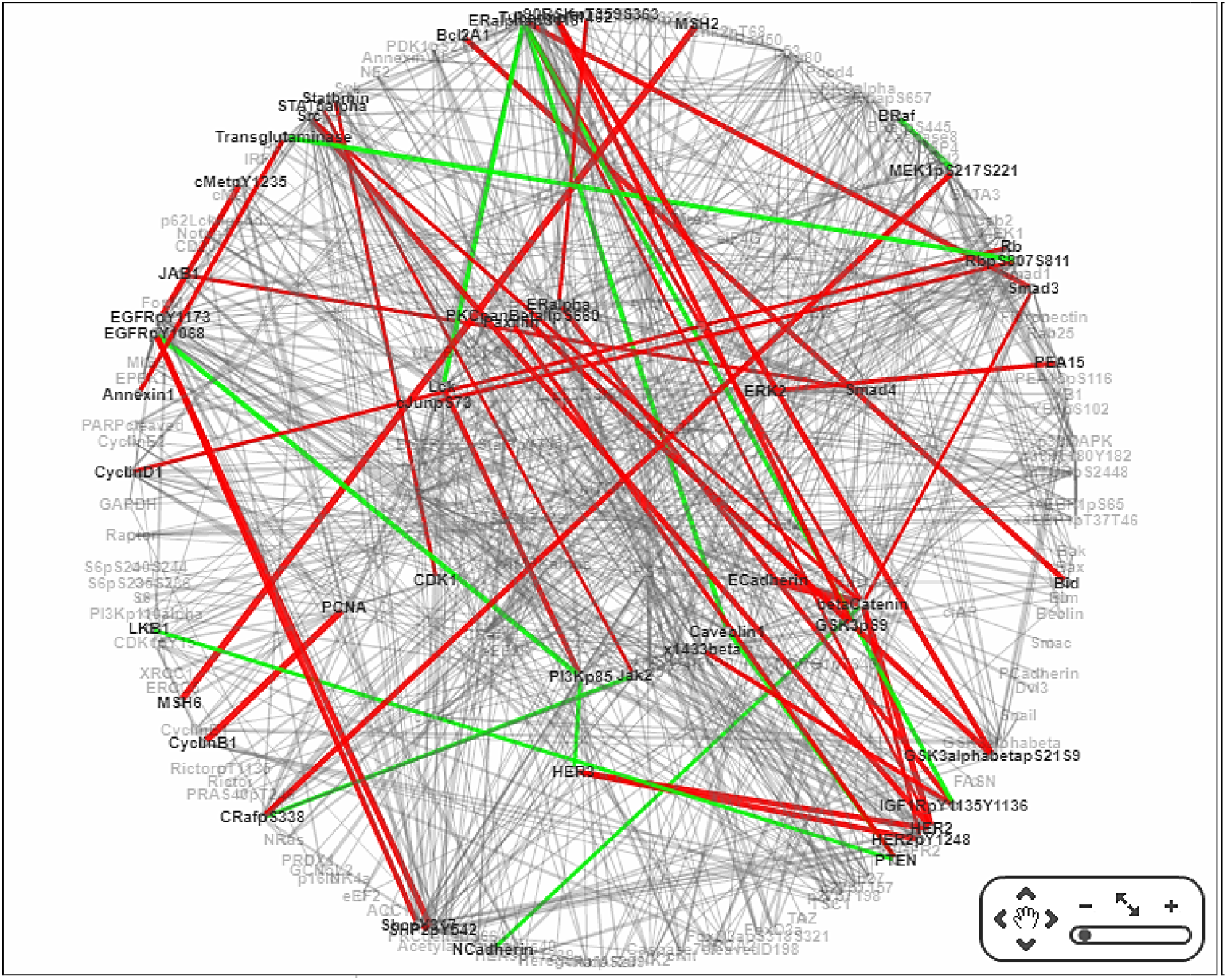
Protein-protein interaction analysis of human bladder cancer samples. Positive correlation between proteins were represented in red lines and negative correlation between proteins were shown in green lines. It is possible to observe several protein interactions that could be applied for new studies involving canine bladder cancer. The visualization was performed using TCPA online tool (https://tcpaportal.org/tcpa/network_visualization.html).

**Table 3.** Proteins in human bladder cancer associated with overall survival in the cohort of 344 patients.

| **Protein** | **Gene** | **Cox P** | **Log-Rank P** |
| --- | --- | --- | --- |
| ANNEXIN1 | ANXA1 | 0.000039282 | 0.0000079633 |
| TAZ | TAZ | 0.012031 | 0.00016332 |
| SRC | SRC | 0.00058508 | 0.00037203 |
| SF2 | SRSF1 | 0.033481 | 0.00073484 |
| ARID1A | ARID1A | 0.021321 | 0.0008096 |
| GATA3 | GATA3 | 0.0015211 | 0.0012084 |
| BAK | BAK1 | 0.068011 | 0.0012196 |
| EGFR | EGFR | 0.006267 | 0.0016964 |
| CD20 | MS4A1 | 0.049073 | 0.0071387 |
| SCD1 | SCD1 | 0.10015 | 0.0078273 |
| CABL | ABL | 0.028914 | 0.011089 |
| GATA6 | GATA6 | 0.18358 | 0.014776 |
| BAP1C4 | BAP1 | 0.41571 | 0.015214 |
| RICTOR | RICTOR | 0.010638 | 0.015853 |
| AXL | AXL | 0.15225 | 0.015863 |
| SMAD3 | SMAD3 | 0.0048029 | 0.017136 |
| BRAF_pS445 | BRAF | 0.068071 | 0.017645 |
| SMAC | DIABLO | 0.0039151 | 0.017688 |
| BECLIN | BECN1 | 0.027124 | 0.018469 |
| PARPCLEAVED | PARP1 | 0.21592 | 0.018488 |
| ADAR1 | ADAR | 0.050259 | 0.022731 |
| SHC_pY317 | SHC1 | 0.23036 | 0.027714 |
| TRANSGLUTAMINASE | TGM2 | 0.53787 | 0.036741 |
| ANNEXINVII | ANXA7 | 0.36004 | 0.045156 |
| PEA15 | PEA15 | 0.15168 | 0.046395 |
| PKCALPHA | PRKCA | 0.056178 | 0.047571 |
| MEK1 | MAP2K1 | 0.65034 | 0.051509 |
| CAVEOLIN1 | CAV1 | 0.03709 | 0.058778 |

**Figure 5.**
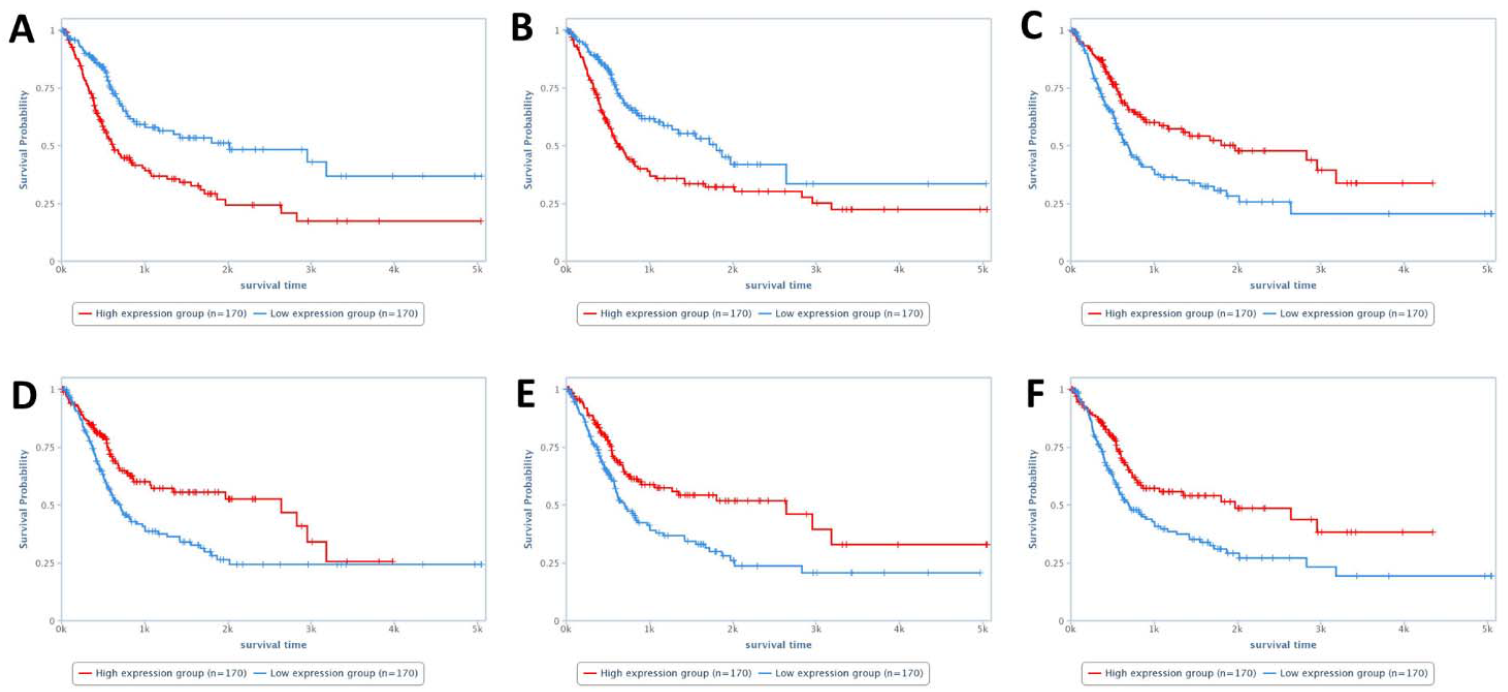
Survival analysis of the human patients with bladder cancer. A: patients presenting high Annexin1 expression experienced a shorter survival time. B: overall survival of patients according to TAZ expression. Patients with high TAZ expression experienced a shorter survival time. C: overall survival according to SRC expression. Patients with lower SRC expression experienced a shorter survival time. D, E and F: overall survival according to SF2, ARID1A and GATA3 expression, respectively. For these three proteins, patients with lower expression experienced a shorter survival time. Survival analysis performed using TCPA online tool (https://tcpaportal.org/tcpa/survival_analysis.html).

## 4 Discussion

The human bladder cancer molecular knowledge it is widely described in the literature and the deposit of these previous publish data brings the opportunity to reanalyze this data, providing new insights for comparative oncology. Although dogs can be considered models to human bladder cancer, few studies provided a full description of the canine bladder cancer molecular data. Since most canine bladder cancer studies have published isolated assessments of different proteins, the present study extracted these data and evaluated them together, to understand how these proteins could interact with each other and identify a profile of the previous veterinary studies.

In the meta-analysis, after the first evaluation, a high number of studies were excluded because they had no concomitant normal tissue analysis (N=31/56). The inclusion of normal tissue is important to stablish a pattern of expression between normal and cancer tissue (Kosti et al., 2016). During the carcinogenic process, cancer cells can change their expression profile with gains or losses of expression of several genes, being important to include normal samples to avoid bias (Kosti et al., 2016). Besides that, for some in-silico analysis, it is necessary to have a p-value related to the protein expression in tumor cell compared to normal tissue.

The meta-analysis demonstrated that most of veterinary studies did not evaluate muscle invasion (23/25) by the tumor or did not provide a clear information regarding this topic. Thus, as a future direction for canine studies evaluating transitional cell carcinoma from the bladder, we strongly suggest authors to evaluate muscle-invasion to provide a stronger evidence regarding dogs being models to human bladder cancer.

Most of the published manuscript that met inclusion criteria, evaluated one up to three proteins or genes and only five previous manuscript performed large-scale analysis. We extracted protein or gene information from manuscript with isolated proteins and evaluated these proteins together to identify the profile of these previous studies. All studies with isolated proteins were focused on the evaluation of oncogenes. Interestingly, most of them evaluated tyrosine kinase receptors, such as ERBB2, EGFR, VEGFR and PDGFR (Tsuboi et al. 2019, Walters et al. 2017, Hanazono et al. 2014). Our PPI analysis revealed a high number of interactions among these proteins, even though they have been evaluated separately in each study. Since they present these high interaction level, future studies can use our PPI analysis to select connected proteins to evaluate its prognostic or predictive value in canine bladder cancer. Interestingly, the ontology analysis of the studies with isolated proteins revealed several terms related with tyrosine kinase activity, phosphorylation process and alteration of ERK1 and ERK2 cascade. Thus, the previous literature was focused on the search of small molecules inhibitors targets. On the other hand, until now, no small molecules inhibitors have been successful proposed in the treatment of canine bladder cancer.

The canine transcriptome data cross validated with TCGA data, revealed 35 genes with high probability to present deregulation in both human and canine bladder cancer. Since these data were obtained from five different canine studies and the 344 human samples from TCGA, it seems consistent and could be used for further investigation. Once it can be difficult to identify a potential marker to be tested in a future study, our list provides markers with strong potential that are up or down regulated in both human and canine bladder cancer. One important limitation of our analysis was the absence of muscle invasion information/standardization in canine samples. Thus, we can lack important genes related to muscle-invasiveness as a known worst prognosis finding. Nevertheless, choosing a gene for future studies from this list could have a higher potential than a random search. Besides that, we did an analysis on the data from 344 human bladder cancer to identify the genes related to patient’s overall survival. Since in veterinary medicine the survival data is usually absent in the published studies, considering human survival data is a unique opportunity to identify candidates related with prognostic potential in veterinary oncology.

Among the 28 proteins with prognostic value, we identified Annexin1, GATA-3 and EGFR. Annexin1 overexpression was previously associated with tumor progression and was considered an independent marker for metastasis-free survival prediction (Li et al., 2010). Besides that, Annexin1 expression was also previously associated with chemotherapy relapse and resistance in human bladder cancer (Yu et al., 2014). In the present meta-analysis, studies evaluating e valuating Annexin1 expression in canine bladder tumors were not founded. GATA3 is widely used in human medicine as a diagnostic marker (Inoue et al., 2017, Agarwal et al. 2019). Interestingly, besides a diagnostic marker, GATA3 has been shown as an important prognostic marker in human bladder cancer (Inoue et al., 2017). GATA3 decreased expression is associated with lower recurrence-free survival, high frequency of muscle invasiveness and positive correlation with tumor progression. In veterinary medicine, one previous review manuscript mentioned GATA3 expression in canine bladder cancer and showed a figure of GATA3 expression in a sample of canine bladder carcinoma (Knapp et al., 2014). However, since it was a review manuscript, these authors did not evaluate canine bladder carcinoma samples. The corresponding author was contacted and kindly provided information regarding the GATA3 antibody used in the case of the image. Thus, GATA3 has a potential to be a prognostic marker for canine bladder cancer.

EGFR it is an important marker in human bladder cancer, being associated with overall survival, muscle invasiveness and tumor recurrence (Hashmi et al. 2018). Thus, EGFR overexpression is studied as target for anti-EGFR therapies (Karyagina et al., 2020). In dogs, EGFR was previously evaluated in canine bladder cancer (Hanazono et al., 2015). However, these authors evaluated EGFR gene and protein in samples with no association with clinical pathological findings. However, based on meta and *in-silico* analysis, EGFR shows a promising potential to be both prognostic and predictive value in canine bladder cancer. Nagaya et al. (2018) evaluated an anti-EGFR monoclonal antibody in canine transitional carcinoma cells from bladder *in vitro* and *in vivo*. The authors finding suggested that this anti-EGFR monoclonal antibody could be promising for the treatment of dogs with bladder cancer. Thus, both humans and dogs can benefit from clinical trials involving anti-EGFR in a veterinary model.

## 5 Conclusion

The previous canine literature on bladder cancer was focused on the evaluation of isolated markers with no association with patient’s survival. Besides that, the lack of information regarding tumor muscle-invasion can be considered an important limitation when comparing human and canine bladder tumors. Our in-silico analysis involving canine and human transcriptome data provided several genes with potential to be markers for both human and canine bladder tumors and these genes should be considered for future studies on canine bladder cancer.

## 7 Author Contributions

VVG and CEF-A wrote the first manuscript draft. VVG performed the meta-analysis and in silico analysis of the selected data. CEF-A and RFA checked the meta-analysis and the in-silico data independently. RL-A and VG contributed with constructive comments. CEF-A supervised the project. All authors read and approved the final manuscript.

