## Supplementary Figure 1 for "A comparative meta and *in silico* analysis of differentially expressed genes and proteins in canine and human bladder cancer"

**Supplementary Table 1**. Ontology analysis of the isolated proteins extracted from the previous published papers on canine transitional cell carcinoma. A, B and C represents the ontologies visualization on REVIGO (<http://revigo.irb.hr/>), considering ontology process with statistical difference and excluding redundancies. D, E and F represents ontology analysis on Enrichr (<https://amp.pharm.mssm.edu/Enrichr/>) demonstrating the ontology process by p-value (red bars). A and D are molecular function-related ontologies, B and E, are ontologies related to Cellular component and C and F Biological-process related analysis. This analysis revealed that the published literature is focused on the study of oncogenes with tyrosine-kinase properties. The regulation of tyrosine kinase receptors and phosphorylation were the most common terms.

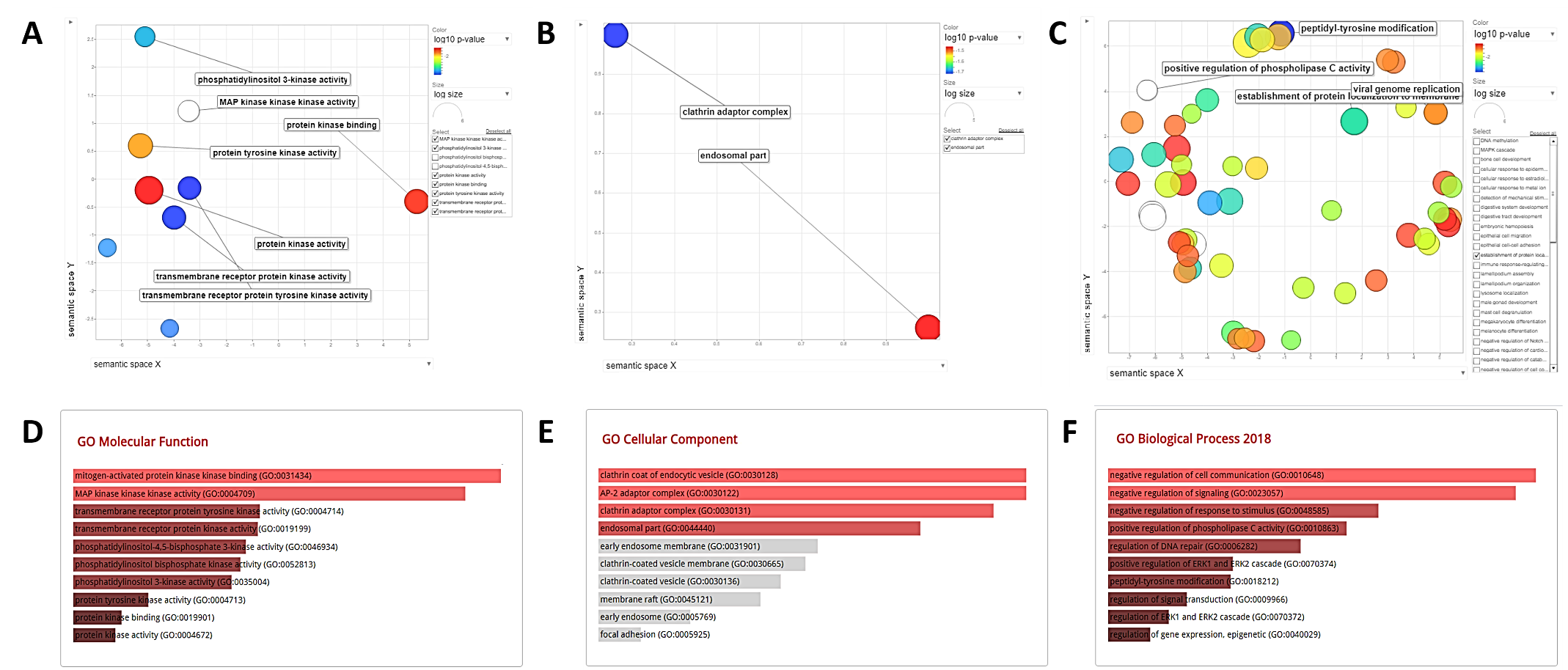
